# The P300 as marker of inhibitory control – fact or fiction?

**DOI:** 10.1101/694216

**Authors:** René J. Huster, Mari S. Messel, Christina Thunberg, Liisa Raud

**Affiliations:** Multimodal Imaging and Cognitive Control Lab, Department of Psychology, University of Oslo, Oslo, Norway; Cognitive and Translational Neuroscience Cluster, Department of Psychology, University of Oslo, Oslo, Norway

**Keywords:** P300, P3, N200, N2, stop signal task, stopping, inhibition, latency

## Abstract

Inhibitory control, i.e., the ability to stop or suppress actions, thoughts, or memories, represents a prevalent and popular concept in basic and clinical neuroscience as well as psychology. At the same time, it is notoriously difficult to study as successful inhibition is characterized by the absence of a continuously quantifiable direct behavioral marker. It has been suggested that the P3 latency, and here especially its onset latency, may serve as neurophysiological marker of inhibitory control as it correlates with the stop signal reaction time (SSRT). The SSRT estimates the average stopping latency, which itself is unobservable since no overt response is elicited in successful stop trials, based on differences in the distribution of go reaction times and the delay of the stop- relative to the go-signal in stop trials.

In a meta-analysis and an independent EEG experiment, we found that correlations between the P3-latency and the SSRT are indeed replicable, but also unspecific. Not only does the SSRT also correlate with the N2-latency, but both P3- and N2-latency measures show similar or even higher correlations with other behavioral parameters such as the go reaction time or stopping accuracy. The missing specificity of P3-SSRT correlations, together with the general pattern of associations, suggests that these manifest effects are driven by underlying latent processes other than inhibition, such as those associated with the speed-accuracy trade-off.

## 1. Introduction

The ability to quickly stop ongoing actions, thoughts, or memories, is considered a hallmark of executive functions or cognitive control. Impaired inhibitory control has consequently been associated with a number of mental disease states, including attention-deficit/hyperactivity disorder (ADHD), obsessive-compulsive disorder, or substance use disorders (e.g., Nigg et al. 2007). The study of inhibition in the motor domain, so-called response inhibition, serves as a proxy for more cognitive domains in which the actual effects of interest, such as the suppression of memories or urges, are notoriously hard to observe (Aron, 2007).

Response inhibition in humans is most commonly studied using the stop signal task (SST) or Go/No-go task (GNGT), both of which putatively probe the rapid suppression of an already initiated and predominant response pattern. In short, in both tasks a go-signal is presented in the majority of trials (e.g., 75%), and participants are instructed to respond as quickly as possible with a button press. In the remaining trials, the no-go or stop signal instructs the participants to withhold the response. While the no-go signal is presented instead of the go-signal in a no-go trial in the GNGT, the stop-signal follows the go-signal with a short delay (the stop signal delay, SSD) in the SST. By systematically varying the SSD such that participants are unsuccessful at stopping in about 50% of the stop trials, the SST allows for the calculation of the stop signal reaction time (SSRT; e.g., Band et al. 2003). The SSRT, often calculated by subtracting the average SSD from the mean response time to go stimuli, provides an estimate of an individual’s speed of the stopping process, and is considered the purest parameter representing inhibitory control capabilities.

A network formed by the right inferior frontal gyurs, the pre-supplementary motor area, and distinct nuclei of the basal ganglia such as the subthalamic nucleus, are believed to implement the stopping process at the neural level (e.g., Aron et al. 2014). While the bulk of research on this inhibitory control network has been done on the stopping of behavioral responses, recent research suggests that it may be domain-general and thus extends its functions also into the more cognitive realm (e.g., Wessel et al., 2016). Much of the work leading to the identification of this network has been done using functional magnetic resonance imaging and transcranial magnetic stimulation (e.g., Cai et al., 2012). Since fMRI suffers from a low temporal resolution though, recent research has shifted more towards the use of electroencephalography (EEG) to better understand the temporal dynamics of inhibitory control (e.g., Huster et al. 2013). The identification of an unequivocal neural signature of an inhibitory control process is of utmost importance. Such a marker would represent a more direct measure of inhibitory control capabilities than the SSRT, which often cannot easily be interpreted since its computation is prone to strategic as well as maladaptive adjustments of behavior (such as reflected in response slowing or go ommissions; e.g., Matzke et al., 2019; Verbruggen et al., 2019; Verbruggen et al., 2013). As such, it would qualify as diagnostic marker in disorders believed to be associated with deficient inhibition. Not least, a valid neural fingerprint of inhibition proper would constitute a meaningful target for neuromodulatory interventions aiming at the augmentation of inhibitory control capabilities.

One event-related potential (ERP) that has repeatedly been suggested as a potential marker of inhibition is the frontocentral P300, or P3a (e.g., Enriquez-Geppert et al., 2010; Huster et al., 2013; Wessel & Aron, 2015). This positive potential, henceforth simply referred to as P3, is consistently found following stimuli that instruct participants not to respond to a stimulus in a context where responding is the prepotent tendency. At the single trial level, it corresponds to increased frontocentral activity in the low theta and delta frequency range with a relatively strong time-or phase-locking (e.g., Huster et al., 2014; Huster et al., 2017).

A consistent finding is that the P3 is increased under conditions conceptually linked to high inhibitory load. Lowered stop-signal probabilities, cue-induced response preparation prior to stop-signal presentation, or faster average response times, for example, are all associated with increased P3 amplitudes (reviewed in Huster et al., 2013). In addition, potential impairments of inhibitory control are often associated with decreased P3 amplitudes, as seen with ADHD for example (e.g., Bekker et al., 2005; Lansbergen et al., 2007). With respect to the timing of the P3, it has been shown that its peak latencies in unsuccessful stop trials are quite unequivocally delayed relative to successful stop trials (e.g., Kok et al., 2004; Ramautar et al., 2004, 2006). The notion that the P3 latency rather than tis amplitude may be better suited to assess inhibitory control has received some support again recently. Wessel et al. (2015) suggested that the onset of the P3, rather than its peak amplitude or peak latency, may be a more specific marker of response inhibition, as the timing of the P3 onset coincides with the SSRT. And indeed, a positive correlation was found such that participants with longer SSRTs also exhibited later P3 onsets.

Whereas these findings altogether point to the possibility that the P3, by its amplitude and/or latency, may serve as a marker of response inhibition, some conceptual issues still need further clarification. First, experimental manipulations of inhibitory load may often be confounded by other cognitive factors. Decreasing the probability of stop trials may well make it more difficult to successfully inhibit a response, but it could also be argued that the relative increase in “novelty” of these stimuli augments their attentional capture. Secondly, the SSRT is merely an indirect estimate of inhibitory speed, because it is computed as a difference measure from go reaction times and SSDs. Isolated assessments of associations between the SSRT and selected neural measures may therefore be misleading, since the relevance of moderating or mediating third variables remains unspecified. Thirdly, muscle activity recorded from response effectors in successful stop trials suggest that the onset of an inhibitory influence can be seen at around 150 ms post stop-stimulus presentation, thus preceding P3 onset latencies as well as usual SSRT estimates by about 70 ms (Raud & Huster, 2017).

This study set out to investigate the applicability and validity of P3-derived measures as potential markers of response inhibition. We specifically focused on P3 amplitude and latency measures and their relationship with behavioral indices of SST performance. However, to test the specificity of these associations and to study the P3 in its processing context, we also assessed associations of the N2 with behavioral performance measures, as well as the relationship between the N2 and the P3. Here, the N2 refers to a fronto-centrally maximal negative deflection occurring about 200 ms post stimulus presentation. Just as the P3, the N2 is usually larger in stop than in go trials (e.g., Huster et al., 2013). The N2 is believed to be generated in the midcingulate cortex, and to indicate the occurrence of conflicts in information processing, such as response conflict or deviations from expected outcomes of actions (e.g., Huster et al., 2010, 2011, 2012). The first section reports a meta-analysis on studies that specifically tested and reported associations between P3-indices and the SSRT. We then tested hypotheses derived from this meta-analysis on a data set including EEG, electromyography (EMG), and behavioral performance measures of a SST. We focused on the most common quantification methods, i.e. the extraction of peak amplitudes, peak latencies, as well as onset latencies from standard ERPs as well as decomposed EEG, namely component ERPs derived by subject-specific and group-level independent component analysis (SS-ICA and G-ICA, respectively).

## 2. A meta-analysis of P3/N2-SSRT associations

An systematic literature review was conducted to identify published studies that assessed and reported associations of P3-derived measures with the SSRT. We further assessed N2-SSRT associations to check for the functional specificity of the P3 in its processing context. Table 1 lists relevant articles alongside their ERP-parameters and correlations with the SSRT. Dependent measures corresponding to P3 peak or mean amplitude were subsumed under *P3 amp; P3 peak lat* refers to the quantification of the latency at which the P3 showed its maximum amplitude; *P3 onset lat* incorporates variables that quantify the onset of the P3. The same grouping was applied to N2-derived measurements.

**Table 1.** Overview of studies that report correlations between N2-/P3-measures and the SSRT.

| Article | P3 amp | P3 peak lat | P3 onset lat | N2 amp | N2 peak lat | N2 onset lat | N | Comments |
| --- | --- | --- | --- | --- | --- | --- | --- | --- |
| Liottie et al., 2005 | n.s. |  |  |  |  |  | 10/10 | ADHD/HC |
| Johnstone et al., 2007 | n.s. | n.s. |  | n.s. | n.s. |  | 24 |  |
| Van Gaal et al., 2010 | 0.22 |  |  | 0.53 |  |  | 15 |  |
| Anguera et al., 2012 | 0.30 | 0.60 |  | 0.30 | 0.10 |  | 17 | Older adults |
| Hughes et al., 2012 | 0.01/-0.47 | 0.14/-0.35 |  |  |  |  | 10/13 | SZ/HC |
| Senderecka et al., 2012 | -0.51 | 0.29 |  | -0.32 | 0.47 |  | 40 | Pooled across ADHD/HC |
| Jones, et al., 2013 | -0.61/0.43 |  |  |  |  |  | 16 | SSD 150/SSD 150-250 |
| Huster et al., 2014 | 0.61 |  |  |  |  |  | 13 | Exploratory analysis; highest correlation |
| Logemann et al., 2014 | -0.65 |  |  | -- |  |  | 16 | N2 excluded (atypical topography) |
| Wessel et al., 2015 |  | 0.3 | 0.6 |  |  |  | 12/10/14/26 | Pooled across four studies |
| Senderecka et al., 2016 | -0.31 | 0.40 |  | -0.52 | 0.62 |  | 33 |  |
| Wessel et al., 2016 |  |  | 0.36 |  |  |  | 22 |  |
| Raud et al., 2017 | -0.42 |  | -0.28 | 0.16 |  | -0.02 | 30 |  |
| Dutra et al., 2018 |  |  | 0.52 |  |  |  | 17 |  |
| Wessel et al., 2018 |  |  | 0.7 |  |  |  | 18 |  |
| Hoptman et al., 2018 | -0.41 |  |  |  |  |  | 42 | Pooled across SZ/HC |

### 2.1. Procedure

To generate a starting list of articles that may contain P3-SSRT correlations, several searches were conducted in PUBMED with varying keywords, of which the combination of “stop signal task” and “EEG” produced the largest and most encompassing list. We then reviewed every article, selecting those that reported to have assessed associations between P3-derived variables and SSRTs, and adding further articles to the list based on cross-referencing in already reviewed articles. This way, we identified a total of 16 articles that reported tests of P3-SSRT correlations. If studies reported statistically non-significant correlations without specifying the exact correlation coefficient, the authors were contacted, and, upon provision, the correlation coefficient was included in the analysis; if the non-significant effect could not be specified any further, n.s. was entered. For each of the variables, except for the N2 onset latencies, a meta-analysis of the correlations was conducted using the MedCalc software (www.medcalc.org, version 18.9.1). Most variables exhibited a significant degree of heterogeneity, and we therefore calculated the summary correlation coefficient under the random effects model according to DerSimonian and Laird (1986).

### 2.2. Results

Although the table is relatively sparsely beset, the data sufficed for meta-analytic assessment for all ERP-parameters but the N2 onset latency. Figure 1 depicts the weighted study-coefficients, the summary correlation-coefficients, as well as corresponding confidence intervals (CI). With respect to ERP amplitudes, neither the P3 nor the N2 exhibited significant summary correlations with the SSRT (P3 amp: r = −0.21, 95%-CI = −0.44 to 0.04s, p = 0.22; N2 amp: r = −0.01, 95%-CI = −0.39 to 0.37, p = 0.97). In contrast, the three remaining latency measures all revealed significant summary correlations with the SSRT (P3 peak lat: r = 0.3, 95%-CI = 0.09 to 0.49, p < 0.008; P3 onset lat: r = 0.41, 95%-CI = 0.02 to 0.69, p < 0.04; N2 peak lat: r = 0.46, 95%-CI = 0.18 to 0.66, p < 0.003).

**Figure 1.**
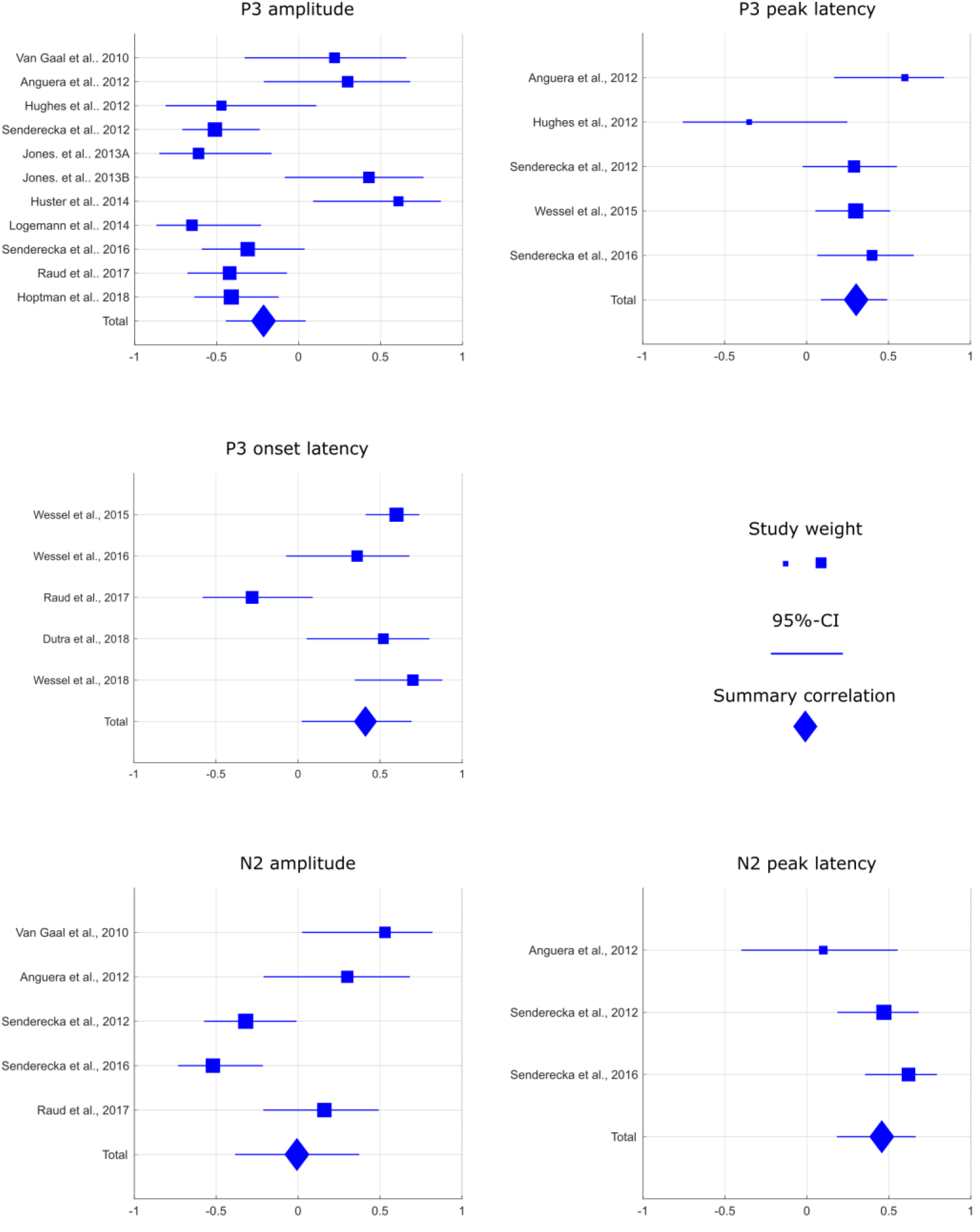
Results of the meta-analysis of correlations between N2-/P3-measures and the SSRT.

### 2.3. Interim Discussion

Current evidence therefore suggests that neither N2 nor P3 amplitudes are consistently correlated with the SSRT. This is somewhat surprising, since a previous review of the EEG literature seems to suggest that conditions of increased inhibitory load coincide with higher P3s, a finding that usually is mirrored when comparing healthy controls and patient groups with potentially impaired inhibitory control.

Latency measures, on the other hand, quite consistently show an association with the SSRT with overall medium effect sizes. Both the P3 peak latency as well as the P3 onset latency are in sum positively correlated with SSRTs, such that earlier P3s coincide with shorter SSRTs. This effect would generally be in accordance with the notion that the latency of the P3 may serve as indicator of inhibitory control capabilities (e.g., Wessel & Aron, 2015). However, this effect is not specific for the P3, since the N2 peak latency shows the same association. For now though, it is unclear whether this is the case for the N2 onset as well (only a single study assessed this association and provided a null finding).

Altogether, this opens the possibility that neither N2 nor P3 latencies serve as specific indicators of the temporal dynamics of inhibition proper, but that these associations may rather be driven by the timing of earlier processing stages (e.g., sensory processing) or by general capacity limitations for higher order cognitive processing.

Since only relatively few studies report relevant correlations, we were unable to assess the relevance of potential moderator variables. Amplitude measures, for example, can be derived in different ways, e.g. by extracting the peak amplitude or by computing the mean amplitude over larger time frames. The same holds true for latency measures; the P3 onset latency, for example, has been quantified based on the earliest significant difference between go and stop trials in some studies, whereas others may choose to compute it as the time point by which a certain percentage of the peak amplitude is reached. Also, whereas the majority of studies focusing on the electrophysiology of stopping relies on parameters directly derived from scalp EEG, more recent studies often apply data decomposition techniques to better isolate the latent processes underlying specific EEG components, e.g. via principal or independent component analysis (PCA and ICA, respectively).

We hope that future studies more regularly assess and report associations of various EEG-derived and behavioral performance measures, so that more sophisticated meta-analyses can be conducted that include the assessment of the effects of potential moderator variables as those mentioned above. Such brain-behavior associations are worth to be assessed and reported, even though it might be in a less specific, or broader and more exploratory manner, as to build a good foundation for summary assessments such as the meta-analyses conducted here. With this in mind, we now proceed to the empirical assessment of ERP-behavior associations.

## 3. A comparative analysis of brain-behavior correlations of stopping

In accordance with our meta-analysis, we now set out to further assess the association of EEG-derived variables and behavioral performance measures as observed in the SST. We will do so predominantly using exploratory analyses. Nonetheless, based on our previous meta-analytic results we can formulate the following expectations: 1) P3 and N2 latency (but not amplitude) measures correlate with the SSRT; 2) since both ERPs show these latency-SSRT associations, suggesting that earlier processing stages may drive these effects, we expect N2 and P3 latencies to be correlated.

To guide our exploratory interpretation of the correlation coefficients we will focus on correlations of |0.2| and above for two reasons: 1) it follows our expectations of the medium effect sizes and their corresponding variation across studies found in the meta-analyses; 2) it compensates for a potential publication bias that is known to cause an overestimation of actual effect sizes. In contrast to the previous analysis, we now will assess the overall structure of associations between ERP and behavioral variables. This is necessary, since the SSRT is a difference measure (derived from the go-RT and SSD), and up to now there is no data that would clarify how the N2/P3 relate to go-RT or stopping accuracy.

We therefore extracted peak amplitudes, peak latencies, and onset latencies from EEG. P3 onset latencies were computed in two different ways: 1) the half-amplitude onset latency (1/2 amp. latency), i.e. the earliest time point at which an ERPs amplitude exceeds half of its peak amplitude when moving backwards in time starting at the peak; 2) the differential onset latency (diff. onset latency), i.e., the earliest time point at which go- and stop-trial activity, matched for motor preparation, differs significantly from each other (Wessel & Aron, 2015). Onset latencies were estimated for the P3 only, a) because of high-single trial variability in case of the N2, and b) to follow the analytic pattern set up with the meta-analyses.

We furthermore extracted and compared dependent variables using different data decomposition techniques based on ICA, namely subject-specific (SS-ICA) and group independent component analysis (G-ICA).Whereas there is no direct indication to believe that standard ERPs and ICA-based ERPs would differ dramatically in this specific context (the stop signal task), this notion has not directly been tested yet. These comparative analyses will give us an indication whether findings generalize regardless of major differences in EEG processing. Please refer to Figure 2 for a depiction of EEG and component time-courses, and to Table 2 for an overview of dependent variables.

**Figure 2.**
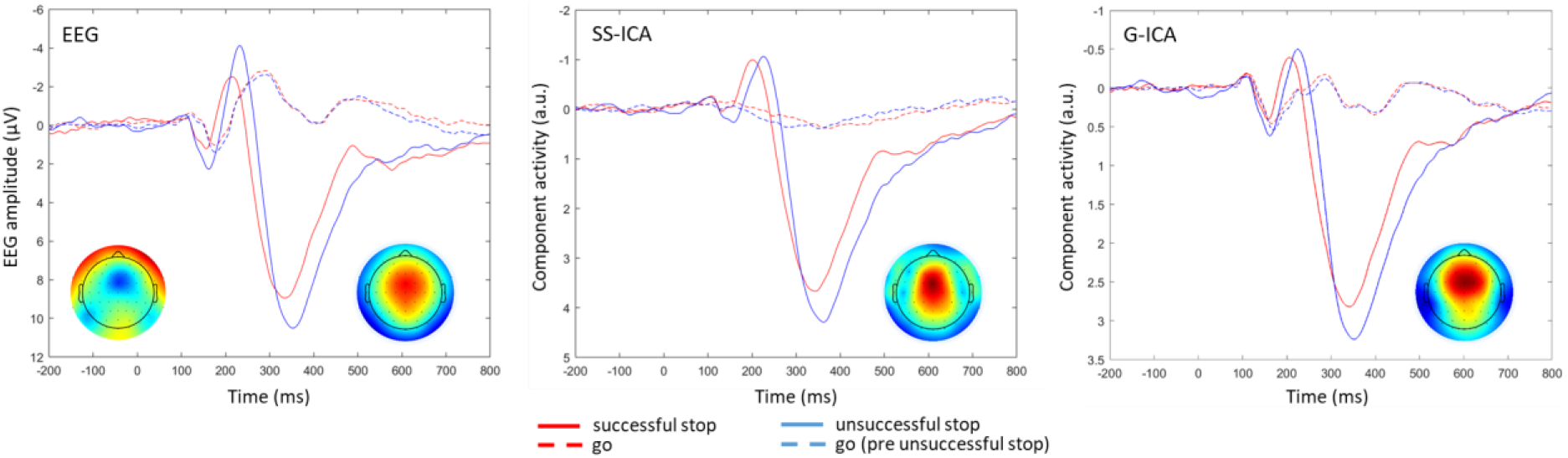
Depiction of the ERPs derived from normal EEG analysis, after subject-specific ICA, as well as group-ICA.

**Table 2.** Descriptive statistics of the dependent measures (means and standard deviations). P3 amplitudes are reported in μV for EEG as computed at electrode Cz. SS-ICA amplitudes for the N2/P3 are not reported because subject-specific ICAs are scaled independently for each subject, and thus not directly comparable. G-ICA amplitudes reflect arbitrary units with common scaling. Latencies are given in ms. The stopping accuracy details the percentage of successful stop trials. ACC = stopping accuracy; EMG = stopping EMG latency.

| Descriptive Statistics |  |  |  |  |  |  |
| --- | --- | --- | --- | --- | --- | --- |
|  | EEG | SS-ICA |  |  | G-ICA |  |
| **P3 amplitude** | 8.61 +/- 3.46 | -- |  |  | 2.95 +/- 1.76 |  |
| **P3 peak latency** | 353 +/- 34.58 | 343 +/- 31.89 |  |  | 343 +/- 25.69 |  |
| **P3 ½ amp. onset latency** | 279 +/- 23.68 | 284 +/- 27.24 |  |  | 278 +/- 21.46 |  |
| **P3 diff. onset latency** | 258 +/- 22.24 | 276 +/- 37.82 |  |  | 262 +/- 32.93 |  |
| **N2 amplitude** | -2.49 +/- 2.08 | -- |  |  | -0.57 +/- 0.81 |  |
| **N2 peak latency** | 212 +/- 20.26 | 209 +/- 29.61 |  |  | 206 +/- 21.28 |  |
|  | BEHAVIOR | Go-RT | FA-RT | SSRT | ACC | EMG |
| **Go-RT** | 605 +/- 89.01 | - |  |  |  |  |
| **FA-RT** | 531 +/- 85.67 | 0.98 | - |  |  |  |
| **SSRT** | 189 +/- 35.48 | -0.38 | -0.40 | - |  |  |
| **Stopping accuracy** | 52.26 +/- 2.41 | 0.88 | 0.88 | -0.54 | - |  |
| **Stopping EMG latency** | 135 +/- 17.28 | -0.55 | -0.49 | 0.36 | -0.48 | - |

### 3.1. Participants

Thirty-seven right-handed participants between the age of 19 and 35 years took part in the study. None reported a history of psychiatric or neurological disorders; all had normal or corrected-to-normal vision. Data from four participants was discarded due to low performance on stop signal task. The final sample consisted of 33 participants (18 female, 15 male; mean age = 26.6 years). All participants gave written informed consent prior to study participation. The study protocol was approved by the institutional review board of the Department of Psychology at the University of Oslo, and followed ethical standards according to the Declaration of Helsinki.

### 3.2. Task

All participants performed both a GNGT and a SST in a single session, of which here only the SST is of relevance. Task order was counterbalanced across subjects. The SST lasted for about 30 minutes. Task presentation was controlled via E-prime 2.0.

Go-stimuli were green arrows that pointed either to the left or to the right (arrowheads of size 3cm x 3.5 cm). Stop stimuli were blue arrows of the same size as the go stimuli. The participants were seated at a viewing distance of approximately 80 cm from the screen. All stimuli were presented at the center of the screen. If none of the target stimuli was presented, a fixation cross was presented instead at the same location.

Participants were instructed to respond as quickly as possible via a button press with the thumb of the hand corresponding to the direction of the go-stimulus. In stop trials, the stop-stimulus appeared after the go-stimulus with a short delay (the stop-signal delay; SSD), instructing the participant to suppress their already initiated response. Stimuli were presented for 100 ms and the SSD adapted according to a tracking procedure aiming at a stopping accuracy of 50%. After successful stop-trials, the SSD was increased by 50 ms, whereas it was decreased by the same time after unsuccessful stop trials. The minimum and maximum SSD were set to 100 and 800 ms, respectively. The SSD tracking was done separately for the left and right hand. The inter-trial interval was randomly varied between1500-2500 ms.

The task consisted of 800 trials, of which 600 were go trials and 200 stop trials, with an equal number of left- and right-hand trials. Blocks of 80 trials were followed by a feedback that instructed participants to respond faster if the average go reaction time of the preceding block exceeded 500 ms. Instantaneous feedback (“Too slow!”) was given after a go omission or if the reaction time exceeded 100 ms. Prior to the SST, participants completed a short training session of 20 trials. It was stressed that it was not possible to be correct all the time and that it was important to be both fast and accurate.

### 3.3. Data Acquisition

EEG and EMG were recorded using a Neuroscan SynAmps2 amplifier with a sampling rate of 2500 Hz, an online high-pass filter at 0.15 Hz, and an online low-pass filter at 1000 Hz. EEG was measured from 64 passive AG/Ag-Cl electrodes placed in accordance with the extended 10-20 system with two additional horizontal EOG channels placed beside the left and the right eye. All EEG electrodes were referenced online against a nose-tip electrode. Impedances were kept under 5 kOhm. For the EMG, the same type of Ag/Ag-Cl electrodes were used in bipolar recording schemes with placements above the abductor pollicis brevis. The ground electrode was placed on the left arm. The participants’ arms were supported using pillows to reduce spurious baseline muscle tension.

### 3.4. Analyses

Go trial reaction times, stopping accuracies, unsuccessful stop trial reaction times, as well as the SSRT were computed. The SSRTs were estimated separately for left and right hand responses using the integration method, i.e. by subtracting the mean SSD from the go-reaction time distribution percentile corresponding to the probability of unsuccessful stopping. All behavioral measures will be reported after averaging across both hands.

Another “behavioral” index of stopping, the partial response EMG (prEMG) activity in successful stop trials, was derived from the EMG recordings. EMG channels were filtered between 10-200 Hz, resampled to 500 Hz, and segmented relative to the stop stimulus. Trials with amplifier saturation were discarded from the analysis. After computing the root mean square for each time point, a moving average with a window width of 11 data points was applied. The time-series of each trial was then transformed through division by the trial-specific average of pre-go activity from −200 to 0 ms. The single-trial data were then z-scored across all trials and time-points, separately for each hand. New data segments were then extracted from −600 to 1000 ms relative to stop-stimulus in successful stop trials, and these trials were then baseline-corrected by subtracting the average trial-specific baseline from −600 to −400 ms from the whole time-series. This procedure was established to correct for differences basic muscle tension. An automatic algorithm was used to detect EMG bursts, defined as z-scored and baseline corrected activity that exceeded the threshold of 1.2. Then, the prEMG peak latency was calculated for the average of all successful stop trials.

EEG channels were filtered between 0.1 and 80 Hz, resampled to 500 Hz, and re-referenced to the common average reference computed over all EEG channels. Infomax independent component analysis (ICA) was run on the data using the routines provided in EEGLAB, and components capturing eye or muscle artifacts were identified and rejected manually. Data were then subjected to another low-pass filter at 40 Hz, and epochs from −200 to 800ms relative to the go- and stop-stimulus were extracted for valid go-, as well as successful and unsuccessful stop-trials, respectively. A baseline-correction was computed using the −200 to 0ms interval and subtracting the baseline average from the whole time series of a trial. An automatic artefact rejection algorithm was run as implemented in EEGLAB’s pop_autorej-function, and the remaining epochs were visually inspected for residual artifacts to correct those manually.

To identify the P3-component based on subject-specific ICA (SS-ICA), an ICA was run on the cleaned data with the number of extracted components equal to the number of EEG electrodes minus the number of components rejected during artifact correction for a given data set. The resulting components were then inspected and the component capturing the P300 (based on topography and time course) was selected for further processing.

To compute a group independent component analysis (G-ICA), 100 go- as well as 60 successful and 40 unsuccessful stop-trials were randomly selected from each data set (with equal contribution of left- and right-hand trials). These numbers were based on the minimum amount of trials available across data sets, while still allowing for the calculation of reliable ERPs, since the organization of G-ICA relies on the concatenation of equally-sized and structured data sets. G-ICA concurrently estimates a component structure representative for the whole group of data sets by combining subject-specific and group-level principal component analysis (PCA) with a group-level ICA (for details, please refer to Eichele et al., 2011; or Huster et al., 2015, 2018). Here, we extracted a total of 8 components, since the first-level PCA indicated that 8 components explained about 90% variance in each of the single-subject data sets. We then identified the group-component that captured the P300 (based on the component time course and topography). This component was reconstructed for each single data set by extracting the subject-specific demixing matrix and applying it to the single-trial EEG data of each subject, thereby reconstructing all available go- and stop-trials at the component level (a more detailed description as well as example code can be found in Huster et al., 2018).

Event-related potentials were computed for correct go-, as well as successful and unsuccessful stop trials for i) the normal EEG data, ii) the subject-specific P3-components extracted using SS-ICA, and iii) the P3-components reconstructed for each subject using G-ICA. Thus, these ERPs differed in their preprocessing, but otherwise contained the exact same trials. We extracted the N2 peak amplitude and peak latency, as well as the P3 peak amplitude, peak latency, ½-amplitude onset latency, as well as the differential onset latency. The basic time windows were defined as 150-300ms, and 225-420ms for the N2 and P3 measures, respectively. The N2/P3 were defined as the most negative/positive value within respective time windows. The peak amplitude was calculated as the mean amplitude at peak +/- 5 data points, and the peak latency simply as the latency of the local minimum/maximum relative to stop-stimulus onset. The P3 ½-amplitude onset was computed as the time point at which the amplitude first reached a value smaller than peak amplitude when tracing amplitudes backward in time starting at peak. To compute the P3 diff. onset latency, go- and stop trials of the same SSD were matched to control for the influence of motor preparation, and then permutation based statistics were used to determine the earliest time point at which go- and stop-trial activity differed significantly from each other (for details please refer to Wessel & Aron, 2015). Single-trial N2 and P3 amplitudes were computed by extracting for each trial the mean amplitude of a 100ms time window centered around each individual’s N2 or P3 peak latency.

Correlation coefficients were computed as standard bivariate product moment correlations. The correlations between the dependent variables were then visualized by means of graph construction. First, simple graphs were computed for each of the data processing methods by setting a specific threshold for the correlations. Correlations exceeding this threshold (as |r|) contributed edges to the graph, whereas the behavioral or EEG variables constituted the nodes. We then integrated the structures of the graphs derived for each of the three methods by computing a multigraph that thus could contain up to three edges between each pair of nodes. At last, a simple graph was computed by removing the redundant edges of the multigraph. This procedure was repeated with four different thresholds (r = 0.2, 0.3, 0.4, and 0.5) to highlight the graph structure and its change based on effect size.

### 3.5. Results

#### 3.5.1. Behavioral data

Overall, behavioral performance measures were within the normal range of what would be expected of a plain visual stop signal task with an average goRT of 605 ms, an SSRT of 189 ms, and a stopping accuracy of 52%. RTs for unsuccessful stop trials (531 ms) were significantly shorter than normal goRTs (t_32_ = −22.05, p < 0.001); this was also the case for every single participant. The behavioral performance measures also exhibited substantial inter-correlations, which are listed in Table 2 together with other descriptive statistics.

#### 3.5.2. EEG and decomposed ERPs

Figure 2 depicts the ERP and component time-courses and topographies reflecting the N2 and P3. The depicted N2/P3-complex is most pronounced at fronto-central areas of the scalp. Relative to the EEG-ERPs, both ICA procedures seem to dissociate the N2/P3-complex from other EEG phenomena. This can, for example, be seen with the N2, which shows some spatio-temporal overlap with activity over occipital areas, or with the slight drift in the baseline EEG of stop trials. This dissociation of different sources is also reflected in nominally higher mean SNRs for the ICA-procedures as compared to standard EEG processing (EEG: 15.13; SS-ICA: 19.83; G-ICA: 20.21).

It was also tested whether the latencies and amplitudes of the N2 and P3 differed between successful and unsuccessful stop trials, because, at least for the P3, this is commonly reported in the literature. Indeed, P3 peak amplitudes were larger for unsuccessful stop trials with all three analysis methods (EEG: t_32_ = −3.34, p < 0.01; SS-ICA: t_32_ = −3.89, p < 0.001; G-ICA: t_32_ = −4.34, p < 0.001), and P3 peak latencies were significantly later (EEG: t_32_ = −6.98, p < 0.001; SS-ICA: t_32_ = −4.35, p < 0.001; G-ICA: t_32_ = −3.78, p < 0.001). N2 amplitudes were significantly larger (more negative) with EEG (t_32_ = 3.41, p < 0.01), but not with SS-ICA (t_32_ = 1.14, p = 0.26) or G-ICA (t_32_ = 1.58, p = 0.12). N2 peak latencies were also delayed in unsuccessful relative to successful stop trials (EEG: t_32_ = −3.2, p < 0.01; SS-ICA: t_32_ = −4.05, p < 0.001; G-ICA: t_32_ = −3.25, p < 0.01).

#### 3.5.3. P3/N2-SSRT associations

The data indicated that both N2 and P3 latencies are associated with the SSRT. Please refer to Table 3 for the exact correlation coefficients. P3 peak and onset latencies obtained from EEG and ICA-decompositions overall showed medium-sized correlations with the SSRT in the expected direction, such that later P3 peaks were associated with longer SSRTs. The same pattern emerged for the N2 peak latency (except for the G-ICA-based latency measure), with shorter latencies corresponding to shorter SSRTs.

**Table 3.** Correlation coefficients between the ERP-derived latency and amplitude measures and the SSRT.

| SSRT correlations |  |  |  |
| --- | --- | --- | --- |
|  | EEG | SS-ICA | G-ICA |
| P3 amplitude | -0.19 | -- | -0.02 |
| P3 peak latency | 0.19 | 0.30 | 0.24 |
| P3 ½ amp. onset latency | 0.29 | 0.25 | 0.25 |
| P3 diff. onset latency | 0.22 | 0.12 | 0.25 |
| N2 amplitude | -0.27 | -- | -0.08 |
| N2 peak latency | 0.26 | 0.21 | 0.02 |

P3 amplitudes did not correlate highly with the SSRT; EEG-derived N2 amplitudes exhibited a small-to-medium negative correlation though: larger (i.e., more negative) ERPs were associated with longer SSRTs.

N2 and P3 peak latencies exhibited relevant positive correlations with each other with r = 0.39 for EEG, r = 0.55 for SS-ICA, and r = 0.26 for G-ICA. Similarly, N2 and P3 amplitudes were negatively correlated when extracted via SS-ICA (r = −0.49) and G-ICA (r = −0.63), such that more negative going N2s co-occurred with larger P3s. For EEG-derived amplitude measures, the correlation was found to be r = 0.02 only.

#### 3.5.4. Exploratory P3/N2-behavior correlations

We conducted further exploratory correlational analyses between the ERP-derived amplitude and latency measures on the one hand, and goRT, stopping accuracy, and residual EMG activity in successful stop trials on the other hand. Most studies assess or report correlations rather selectively, usually focusing on the SSRT. However, since the SSRT is a difference measure derived from goRTs and SSDs, it is important to also inspect the overall correlation structure. Table 4 lists the correlation coefficients.

**Table 4.** Exploratory correlations of ERP-derived measures with goRT, stopping accuracy, and stopping EMG. Correlations larger than 0.3 were considered relevant and highlighted in bold/italic.

| **goRT** |  |  |  |
| --- | --- | --- | --- |
|  | **EEG** | **SS-ICA** | **G-ICA** |
| **P3 amplitude** | 0.48 | -- | 0.47 |
| **P3 peak latency** | -0.14 | -0.20 | -0.21 |
| **P3 ½ amp. onset latency** | -0.34 | -0.22 | -0.08 |
| **P3 diff. onset latency** | -0.50 | -0.32 | -0.37 |
| **N2 amplitude** | 0.12 | -- | -0.36 |
| **N2 peak latency** | -0.42 | -0.51 | -0.11 |

| **Stopping accuracy** |  |  |  |
| --- | --- | --- | --- |
|  | **EEG** | **SS-ICA** | **G-ICA** |
| **P3 amplitude** | 0.37 | -- | 0.38 |
| **P3 peak latency** | -0.17 | -0.23 | -0.20 |
| **P3 ½ amp. onset latency** | -0.31 | -0.20 | -0.09 |
| **P3 diff. onset latency** | -0.50 | -0.32 | -0.46 |
| **N2 amplitude** | 0.13 | -- | -0.28 |
| **N2 peak latency** | -0.32 | -0.47 | -0.09 |

| **stopping EMG** |  |  |  |
| --- | --- | --- | --- |
|  | **EEG** | **SS-ICA** | **G-ICA** |
| **P3 amplitude** | -0.17 | -- | -0.24 |
| **P3 peak latency** | 0.41 | 0.54 | 0.40 |
| **P3 ½ amp. onset latency** | 0.46 | 0.36 | 0.29 |
| **P3 diff. onset latency** | 0.26 | 0.31 | 0.36 |
| **N2 amplitude** | -0.05 | -- | 0.22 |
| **N2 peak latency** | 0.64 | 0.51 | 0.08 |

Overall, these exploratory analyses indicated that P3 amplitudes and P3 onset latencies are related to both the average goRT and stopping accuracy. Larger P3 amplitudes and earlier onset latencies co-occurred with longer goRTs and higher stopping accuracy. These effects were, however, less pronounced with the ICA-based ½-amplitude latency measures.

Similar effects were found for the N2. The N2 peak latency was associated with goRTs and stopping accuracies, at least when extracted via EEG or SS-ICA, and N2 amplitudes derived via G-ICA correlated negatively with goRTs. As with the P3, earlier N2 latencies and larger N2 amplitudes (i.e., more negative) were associated with longer goRTs and higher stopping accuracies.

With respect to the peak latency of the prEMG activity in successful stop trials, N2 and P3 latency measures largely exhibited positive correlations. Thus, later peak latencies of residual EMG activity in successful stop trials were associated with later N2 and P3 latencies.

#### 3.5.5. Graph estimation and visualization

Simple graphs were computed to assess the structure of the decencies between the behavioral and EEG-derived variables across the different data processing methods. Specifically, we first integrated the correlational structures of each of the three methods by computing a multigraph based on a specified threshold. Correlations exceeding this threshold (as |r|) contributed edges to the graph, whereas the behavioral or EEG variables constituted the nodes. From these multigraphs (i.e., up to three edges between two nodes were possible), a simple graph was computed by removing redundant edges. This procedure was repeated with four different thresholds (r = 0.2, 0.3, 0.4, and 0.5) to highlight the graph structure and its change based on effect size. Figure 3 depicts the resulting graphs. Although it could be expected that the variables show clusters according to their modality (behavior vs. EEG), it is interesting to note that the EEG-derived variables further break up into two clusters with medium to high correlations coefficients, namely those quantifying amplitudes and those specifying latencies. It further seems that these clusters show differential associations with behavioral markers. Whereas the amplitude measures (especially P3amp) are predominantly associated with reaction time measures (faRT, goRT), the EEG-latency measures (especially N2 and P3 peak latencies) exhibit stronger associations with the prEMG latency, the stopping accuracies, as well as the goRT. Overall, associations with the SSRT are weaker than those aforementioned, suggesting that correlations between EEG-derived variables and the SSRT may be mediated through other behavioral variables.

**Figure 3.**
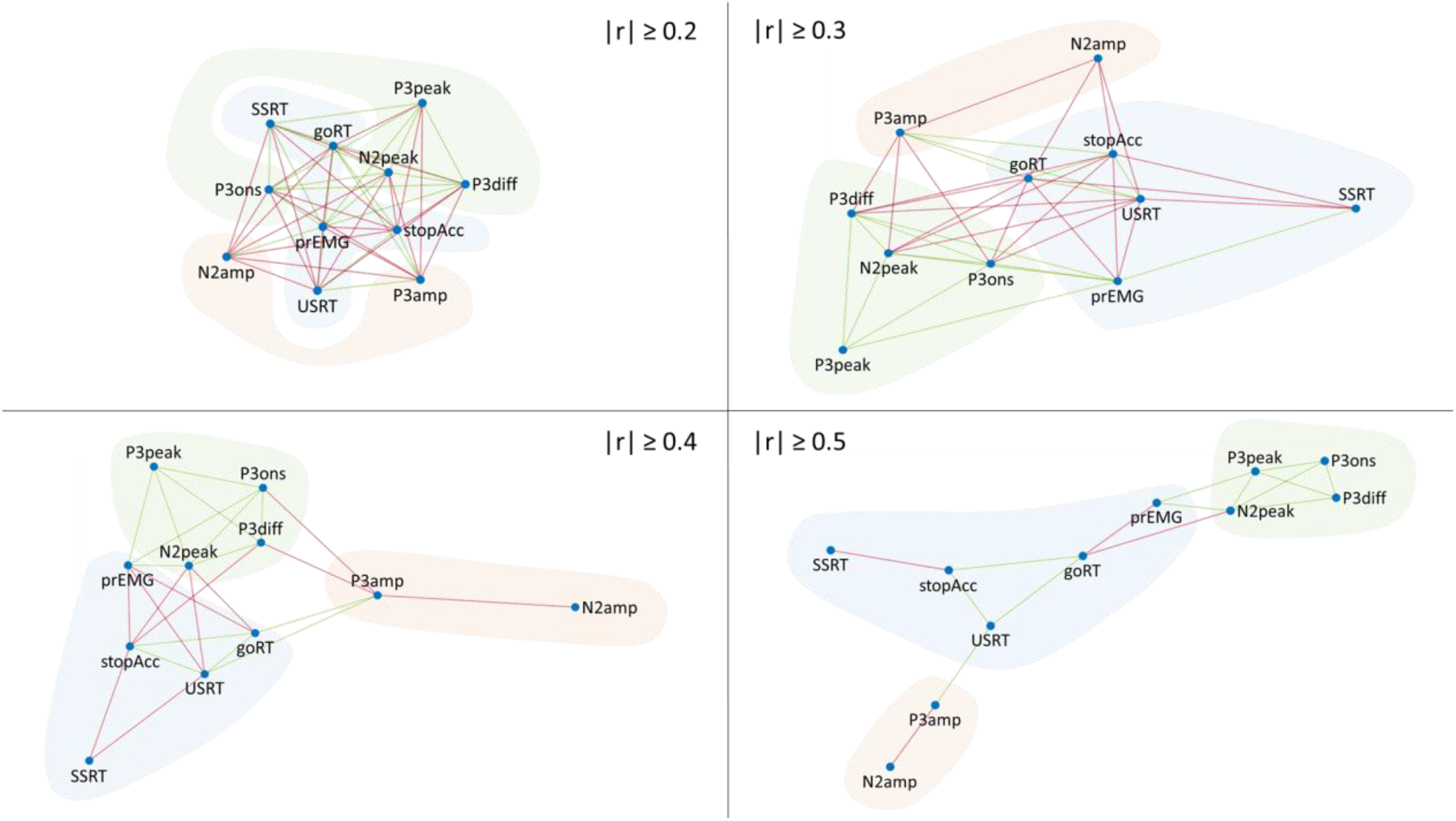
Simple graphs based on the integration of the correlational dependencies derived for EEG, SS-ICA, and G-ICA. Four graphs were established based on the thresholds depicted specified in the figure. Colored clouds visually group the dependent variables: blue – behavioral variables, red – EEG amplitudes, green – EEG latencies. Greed edges denote positive, red edges negative correlations.

## 4. Discussion

Both the meta-analytic as well as the empirical data replicate the previously reported associations between P3 latency and the SSRT with small to medium effect sizes. However, the overall pattern of correlations suggests that the P3-SSRT correlation is not specific, neither with respect to its underlying EEG phenomenon, the P3 as compared to other ERPs, nor regarding its association with the stopping latency measured.

The meta-analysis conducted here supports the notion that P3 latency measures show associations with the SSRT. However, it also suggests that the same holds true for the N2 peak latency. Neither P3 nor N2 amplitude measures exhibit such associations though, suggesting a certain degree of differential functional specificity of the latency and the amplitude measures. The systematic literature review and the meta-analysis also point to some relevant and serious problems. First, only a small fraction of studies reports to have assessed ERP-SSRT associations, and of those not all report the actual correlation coefficient if statistical testing did not indicate significance. This, and the fact that publication bias probably is prevalent also in the neuroscience literature, suggests that the actual effect size might be smaller than estimated here. Second, the sample sizes of these studies were rather small relative to the observed effects (average N of 22.4 participants). That implies that, even before correcting for potential publication bias, most studies were underpowered: with r = 0.3, α = 0.5, n = 23, and one sided testing, we arrive at a power below 50%. Third, there seems to be a marked tendency for selective testing and/or reporting of effects. Even those studies that specifically aimed at the assessment of brain-behavior associations usually did so by merely testing the effect of interest (e.g., the association of P3 latency and SSRT). Whereas this procedure is commendable in its goal to minimize false positive findings, it comes at the risk of missing potential mediator or moderator variables. After all, the SSRT is a difference measure computed from go-RTs and SSDs.

The empirical analysis of associations of N2 and P3 latency and amplitude measures with behavioral parameters further supports the notion that the P3-SSRT association is less specific than thought. Not only is this association not specific to the P3 onset estimated via an SSD-matching of go and stop trials (e.g., Wessel et al., 2015), but it can similarly be found for the P3 ½-amplitude onset latency as well as the peak latency; even the association with the N2-peak latency is of similar size. More importantly perhaps, it is also not specific to the SSRT, but associations of the ERP latency measures are at least of similar size for the goRT, stopping accuracy and residual EMG. The graph visualization highlights the overall constellation of associations, and the somewhat isolated positioning of the SSRT in the graphs for higher effect sizes seem to suggest that its associations with EEG-derived measures may indeed rather be mediated via other behavioral parameters. The constellation of correlations observed here is compatible with the notion that these ERP-SSRT associations may be driven by individual differences in the speed-accuracy trade-off (SAT). The positive correlations of ERP-latency measures with the SSRT are flanked by (generally higher) negative correlations with the goRT and stopping accuracy. This correlational pattern can be expected given that the behavioral measures are clearly indicative of SAT differences across subjects, with high correlations indicating that subjects exhibiting high stopping accuracy show longer response times and shorter SSRTs. This observation is perfectly in line with experimental evidence showing that also within subjects, the SSRT is lower when experimental conditions motivate correct stopping, and is higher when fast responding is stressed instead (e.g., Leotti & Wager, 2010; Greenhouse and Wessel, 2013). However, if this is indeed the case, and whether further information is hidden in the graph structure (e.g., a differential association of ERP-amplitude and latency measures with response times in unsuccessful stop trials and go trials, respectively) needs further investigation.

Even if we were to accept a certain degree of specificity of P3-latency/SSRT associations, other conceptual problems still remain. Correlations of small to medium size leave the much bigger part of the variation unexplained; with correlations of about 0.3, we achieve variance explanation of less than 10 percent. This seems insufficient to consider P3 latency measures reliable markers of the inhibition latency, or the P3 a “motor inhibition component” (e.g., Dutra et al., 2018). Thus, for both empirical and conceptual reasons, it currently seems ill-posed to consider the P3 or its latency a reliable and valid marker for the timing of inhibition.

Given that many of the associations we find between the P3-derived and behavioral measures also seem to be existent with N2-derived measures, what can we say about the functional specificity of the N2 and P3? Whereas there is sufficient evidence to link the N2 to conflict monitoring or prediction errors, the cognitive process(es) associated with the P3 have been proven to be more evasive (Huster et al., 2013). The P3 has been proposed to reflect many different processes, including inhibition, context updating, novelty processing and attentional orienting, but a definitive functional interpretation has yet to come. Even less is known about the interplay and functional dependence of the N2 and P3. We found that the N2 and P3 were correlated across subjects such that larger (more negative) and later N2s were associated with stronger and later P3s. On the other hand, we also found differential amplitude changes in unsuccessful relative to successful stop trials. Whereas the latencies for both potentials were later for unsuccessful stopping, P3 amplitudes were enlarged for these potentials but N2 amplitudes (after decomposition) were unchanged. Two aspects deserve consideration here. First, smaller P3 amplitudes for unsuccessful compared to successful stop trials have been reported in previous studies (e.g., Greenhouse & Wessel, 2013) and this effect has been one argument to link the P3 to inhibition (with putatively lowered inhibition being associated with stopping failures). However, the results reported here are not unique, because lower P3 amplitudes for successful stops had also been reported earlier, yet seem to have received less attention. Kok et al., (2004), for example, found larger P3s in unsuccessful stop trials, and further reported that the differential association of successful and unsuccessful stop trials with shorter and longer SSDs, respectively, can confound potential P3 amplitude differences between these trial types. Secondly, we found this differential modulation for the P3, but not the N2, further supporting functional specificity of these ERPs even when extracted from the same independent component. In sum, to date it is unclear how the underlying neural and cognitive mechanisms associated with the N2 and P3 interact to shape behavioral performance, and potentially response inhibition.

Lastly, it seems noteworthy that the patterns of associations between neural and behavioral markers were very similar, regardless of whether amplitudes and latency measures were derived directly from the EEG, or from data decomposed via subject-specific our group-level ICA. For the P3-derived measures, relevant associations were of similar size and direction across methods. However, this does not mean that the correspondence is perfect and that these methods can be used interchangeably. This is attested by the higher SNRs of potentials obtained from ICA-procedures, driven by ICA’s well-documented capability to dissociate spatio-temporally overlapping processes. The comparison of component strengths for a specific condition across subjects using SS-ICA is not directly feasible, because of the independent scaling of component time-courses and weights. G-ICA minimizes this problem and thus enables the analysis of component amplitudes across subjects; yet G-ICA has been shown to suffer from decreased sensitivity to source patterns with rather weak time-locking (Huster et al., 2015). The fact that correlations with N2 latencies seemed to be consistently lower with G-ICA compared to the other two procedures may be driven by somewhat poorer phase-locking of the theta-activity that underlies the N2 (e.g., Cohen & Donner, 2013). Thus, whereas the general pattern holds across procedures, additional aspects need to be considered to arrive at an optimal decision regarding the method of choice: when interested in group-comparisons or analyses across subjects, G-ICA seems better suited than SS-ICA, which again might be a better choice for the optimal reconstruction of an individual’s component structure including activity patterns with poor time-locking; standard EEG analyses seem to be the compromise, as they are limited with respect to signal quality, especially when single-trial data are of interest.

### 4.1. Conclusion

The P3 has regularly been implicated in inhibition, not least based on reports that especially its onset latency correlates with the SSRT. Through meta-analyses and empirical data we found this correlation to be replicable, yet also unspecific. Similar correlations can be found for the P3 as well as the N2 peak latency. Even more, correlations of the same ERP indices were higher for other behavioral indices such as the go reaction time and stopping accuracy. It therefore remains to be seen whether the reported associations between the P3 and the SSRT may indeed reflect genuine inhibitory control, or whether they rather result from more general behaviorally adaptive patterns such as speed-accuracy trade-offs.

## Acknowledgements

This work was supported by research funds provided by the Department of Psychology at the University of Oslo. We would further thank Magdalena Senderecka, Matthew Hughes, and Alexander Logemann for providing additional data in support for the meta-analyses.

## References

Anguera, J. A., & Gazzaley, A. (2012). Dissociation of motor and sensory inhibition processes in normal aging. Clinical Neurophysiology: Official Journal of the International Federation of Clinical Neurophysiology, 123(4), 730–740. https://doi.org/10.1016/j.clinph.2011.08.024

Aron, A. R. (2007). The Neural Basis of Inhibition in Cognitive Control. The Neuroscientist, 13(3), 214–228. https://doi.org/10.1177/1073858407299288

Aron, Adam R., Robbins, T. W., & Poldrack, R. A. (2014). Inhibition and the right inferior frontal cortex: One decade on. Trends in Cognitive Sciences, 18(4), 177–185. https://doi.org/10.1016/j.tics.2013.12.003

Band, G. P. H., van der Molen, M. W., & Logan, G. D. (2003). Horse-race model simulations of the stop-signal procedure. Acta Psychologica, 112(2), 105–142. https://doi.org/10.1016/S0001-6918(02)00079-3

Bekker, E. M., Overtoom, C. C. E., Kooij, J. J. S., Buitelaar, J. K., Verbaten, M. N., & Kenemans, J. L. (2005). Disentangling deficits in adults with attention-deficit/hyperactivity disorder. Archives of General Psychiatry, 62(10), 1129–1136. https://doi.org/10.1001/archpsyc.62.10.1129

Cai, W., George, J. S., Verbruggen, F., Chambers, C. D., & Aron, A. R. (2012). The role of the right presupplementary motor area in stopping action: Two studies with event-related transcranial magnetic stimulation. Journal of Neurophysiology, 108(2), 380–389. https://doi.org/10.1152/jn.00132.2012

Cohen, M. X., & Donner, T. H. (2013). Midfrontal conflict-related theta-band power reflects neural oscillations that predict behavior. Journal of Neurophysiology, 110(12), 2752–2763. https://doi.org/10.1152/jn.00479.2013

Dutra, I. C., Waller, D. A., & Wessel, J. R. (2018). Perceptual Surprise Improves Action Stopping by Nonselectively Suppressing Motor Activity via a Neural Mechanism for Motor Inhibition. The Journal of Neuroscience: The Official Journal of the Society for Neuroscience, 38(6), 1482–1492. https://doi.org/10.1523/JNEUROSCI.3091-17.2017

Eichele, T., Rachakonda, S., Brakedal, B., Eikeland, R., & Calhoun, V. D. (2011). EEGIFT: Group independent component analysis for event-related EEG data. Computational Intelligence and Neuroscience, 2011, 129365. https://doi.org/10.1155/2011/129365

Enriquez-Geppert, S., Konrad, C., Pantev, C., & Huster, R. J. (2010). Conflict and inhibition differentially affect the N200/P300 complex in a combined go/nogo and stop-signal task. NeuroImage. https://doi.org/10.1016/j.neuroimage.2010.02.043

Greenhouse, I., & Wessel, J. R. (2013). EEG signatures associated with stopping are sensitive to preparation. Psychophysiology, 50(9), 900–908. https://doi.org/10.1111/psyp.12070

Hoptman, M. J., Parker, E. M., Nair-Collins, S., Dias, E. C., Ross, M. E., DiCostanzo, J. N., … Javitt, D. C. (2018). Sensory and cross-network contributions to response inhibition in patients with schizophrenia. NeuroImage. Clinical, 18, 31–39. https://doi.org/10.1016/j.nicl.2018.01.001

Hughes, M. E., Fulham, W. R., Johnston, P. J., & Michie, P. T. (2012). Stop-signal response inhibition in schizophrenia: Behavioural, event-related potential and functional neuroimaging data. Biological Psychology, 89(1), 220–231. https://doi.org/10.1016/j.biopsycho.2011.10.013

Huster, R J, Eichele, T., Enriquez-Geppert, S., Wollbrink, A., Kugel, H., Konrad, C., & Pantev, C. (2011). Multimodal imaging of functional networks and event-related potentials in performance monitoring. NeuroImage, 56(3), 1588–1597. https://doi.org/10.1016/j.neuroimage.2011.03.039

Huster, R J, Westerhausen, R., Pantev, C., & Konrad, C. (2010). The role of the cingulate cortex as neural generator of the N200 and P300 in a tactile response inhibition task. Human Brain Mapping. https://doi.org/10.1002/hbm.20933

Huster, René J., Enriquez-Geppert, S., Lavallee, C. F., Falkenstein, M., & Herrmann, C. S. (2013). Electroencephalography of response inhibition tasks: Functional networks and cognitive contributions. International Journal of Psychophysiology: Official Journal of the International Organization of Psychophysiology, 87(3), 217–233. https://doi.org/10.1016/j.ijpsycho.2012.08.001

Huster, Rene J, Enriquez-Geppert, S., Pantev, C., & Bruchmann, M. (2012). Variations in midcingulate morphology are related to ERP indices of cognitive control. Brain Structure & Function. https://doi.org/10.1007/s00429-012-0483-5

Huster, Rene J., Plis, S. M., & Calhoun, V. D. (2015). Group-level component analyses of EEG: Validation and evaluation. Frontiers in Neuroscience, 9, 254. https://doi.org/10.3389/fnins.2015.00254

Huster, René J., Plis, S. M., Lavallee, C. F., Calhoun, V. D., & Herrmann, C. S. (2014). Functional and effective connectivity of stopping. NeuroImage, 94, 120–128. https://doi.org/10.1016/j.neuroimage.2014.02.034

Huster, René J., & Raud, L. (2018). A Tutorial Review on Multi-subject Decomposition of EEG. Brain Topography, 31(1), 3–16. https://doi.org/10.1007/s10548-017-0603-x

Huster, René J., Schneider, S., Lavallee, C. F., Enriquez-Geppert, S., & Herrmann, C. S. (2017). Filling the void-enriching the feature space of successful stopping. Human Brain Mapping, 38(3), 1333–1346. https://doi.org/10.1002/hbm.23457

Johnstone, S. J., Dimoska, A., Smith, J. L., Barry, R. J., Pleffer, C. B., Chiswick, D., & Clarke, A. R. (2007). The development of stop-signal and Go/Nogo response inhibition in children aged 7-12 years: Performance and event-related potential indices. International Journal of Psychophysiology: Official Journal of the International Organization of Psychophysiology, 63(1), 25–38. https://doi.org/10.1016/j.ijpsycho.2006.07.001

Jones, A., Field, M., Christiansen, P., & Stancak, A. (2013). P300 during response inhibition is associated with ad-lib alcohol consumption in social drinkers. Journal of Psychopharmacology (Oxford, England), 27(6), 507–514. https://doi.org/10.1177/0269881113485142

Kok, A., Ramautar, J. R., De Ruiter, M. B., Band, G. P. H., & Ridderinkhof, K. R. (2004). ERP components associated with successful and unsuccessful stopping in a stop-signal task. Psychophysiology, 41(1), 9–20. https://doi.org/10.1046/j.1469-8986.2003.00127.x

Lansbergen, M. M., Böcker, K. B. E., Bekker, E. M., & Kenemans, J. L. (2007). Neural correlates of stopping and self-reported impulsivity. Clinical Neurophysiology: Official Journal of the International Federation of Clinical Neurophysiology, 118(9), 2089–2103. https://doi.org/10.1016/j.clinph.2007.06.011

Leotti, L. A., & Wager, T. D. (2010). Motivational influences on response inhibition measures. Journal of Experimental Psychology. Human Perception and Performance, 36(2), 430–447. https://doi.org/10.1037/a0016802

Liotti, M., Pliszka, S. R., Perez, R., Kothmann, D., & Woldorff, M. G. (2005). Abnormal brain activity related to performance monitoring and error detection in children with ADHD. Cortex; a Journal Devoted to the Study of the Nervous System and Behavior, 41(3), 377–388.

Logemann, H. N. A., Böcker, K. B. E., Deschamps, P. K. H., Kemner, C., & Kenemans, J. L. (2014). The effect of enhancing cholinergic neurotransmission by nicotine on EEG indices of inhibition in the human brain. Pharmacology, Biochemistry, and Behavior, 122, 89–96. https://doi.org/10.1016/j.pbb.2014.03.019

Matzke, D., Curley, S., Gong, C. Q., & Heathcote, A. (2019). Inhibiting responses to difficult choices. Journal of Experimental Psychology. General, 148(1), 124–142. https://doi.org/10.1037/xge0000525

Nigg, J. T., Carr, L., Martel, M., & Henderson, J. M. (2007). Concepts of inhibition and developmental psychopathology. In D. S. Gorfein & C. M. MacLeod (Eds.), Inhibition in cognition (pp. 259–277). Washington, DC, US: American Psychological Association.

Ramautar, J. R., Kok, A., & Ridderinkhof, K. R. (2004). Effects of stop-signal probability in the stop-signal paradigm: The N2/P3 complex further validated. Brain and Cognition, 56(2), 234–252. https://doi.org/10.1016/j.bandc.2004.07.002

Ramautar, J. R., Kok, A., & Ridderinkhof, K. R. (2006). Effects of stop-signal modality on the N2/P3 complex elicited in the stop-signal paradigm. Biological Psychology, 72(1), 96–109. https://doi.org/10.1016/j.biopsycho.2005.08.001

Raud, L., & Huster, R. J. (2017). The Temporal Dynamics of Response Inhibition and their Modulation by Cognitive Control. Brain Topography, 30(4), 486–501. https://doi.org/10.1007/s10548-017-0566-y

Senderecka, M. (2016). Threatening visual stimuli influence response inhibition and error monitoring: An event-related potential study. Biological Psychology, 113, 24–36. https://doi.org/10.1016/j.biopsycho.2015.11.003

Senderecka, M., Grabowska, A., Szewczyk, J., Gerc, K., & Chmylak, R. (2012). Response inhibition of children with ADHD in the stop-signal task: An event-related potential study. International Journal of Psychophysiology: Official Journal of the International Organization of Psychophysiology, 85(1), 93–105. https://doi.org/10.1016/j.ijpsycho.2011.05.007

van Gaal, S., Ridderinkhof, K. R., van den Wildenberg, W. P. M., & Lamme, V. A. F. (2009). Dissociating consciousness from inhibitory control: Evidence for unconsciously triggered response inhibition in the stop-signal task. Journal of Experimental Psychology. Human Perception and Performance, 35(4), 1129–1139. https://doi.org/10.1037/a0013551

Verbruggen, F., Aron, A. R., Band, G. P., Beste, C., Bissett, P. G., Brockett, A. T., … Boehler, C. N. (2019). A consensus guide to capturing the ability to inhibit actions and impulsive behaviors in the stop-signal task. ELife, 8. https://doi.org/10.7554/eLife.46323

Verbruggen, F., Chambers, C. D., & Logan, G. D. (2013). Fictitious inhibitory differences: How skewness and slowing distort the estimation of stopping latencies. Psychological Science, 24(3), 352–362. https://doi.org/10.1177/0956797612457390

Wessel, J. R. (2018). Prepotent motor activity and inhibitory control demands in different variants of the go/no-go paradigm. Psychophysiology, 55(3). https://doi.org/10.1111/psyp.12871

Wessel, J. R., & Aron, A. R. (2015). It’s not too late: The onset of the frontocentral P3 indexes successful response inhibition in the stop-signal paradigm. Psychophysiology, 52(4), 472–480. https://doi.org/10.1111/psyp.12374

Wessel, J. R., Jenkinson, N., Brittain, J.-S., Voets, S. H. E. M., Aziz, T. Z., & Aron, A. R. (2016). Surprise disrupts cognition via a fronto-basal ganglia suppressive mechanism. Nature Communications, 7, 11195. https://doi.org/10.1038/ncomms11195

